# Scalable, flexible carbon fiber electrode thread arrays for three-dimensional spatial profiling of neurochemical activity in deep brain structures of rodents

**DOI:** 10.1101/2023.04.15.537033

**Authors:** Mingyi Xia, Busra Nur Agca, Tomoko Yoshida, Jiwon Choi, Usamma Amjad, Kade Bose, Nikol Keren, Shahar Zukerman, Michael J. Cima, Ann M. Graybiel, Helen N. Schwerdt

## Abstract

We developed a flexible “electrode-thread” array for recording dopamine neurochemical activity from a lateral distribution of subcortical targets (up to 16) transverse to the axis of insertion. Ultrathin (∼ 10 µm diameter) carbon fiber (CF) electrode-threads (CFETs) are clustered into a tight bundle to introduce them into the brain from a single entry point. The individual CFETs splay laterally in deep brain tissue during insertion due to their innate flexibility. This spatial redistribution allows navigation of the CFETs towards deep brain targets spreading horizontally from the axis of insertion. Commercial “linear” arrays provide single entry insertion but only allow measurements along the axis of insertion. Horizontally configured neurochemical recording arrays inflict separate penetrations for each individual channel (i.e., electrode). We tested functional performance of our CFET arrays *in vivo* for recording dopamine neurochemical dynamics and for providing lateral spread to multiple distributed sites in the striatum of rats. Spatial spread was further characterized using agar brain phantoms to measure electrode deflection as a function of insertion depth. We also developed protocols to slice the embedded CFETs within fixed brain tissue using standard histology techniques. This method allowed extraction of the precise spatial coordinates of the implanted CFETs and their recording sites as integrated with immunohistochemical staining for surrounding anatomical, cytological, and protein expression labels. Neurochemical recording operations tested here can be integrated with already widely established capabilities of CF-based electrodes to record single neuron activity and local field potentials, to enable multi-modal recording functions. Our CFET array has the potential to unlock a wide range of applications, from uncovering the role of neuromodulators in synaptic plasticity, to addressing critical safety barriers in clinical translation towards diagnostic and adaptive treatment in Parkinson’s disease and major mood disorders.

## Introduction

Methods to measure neurochemical activity in the brain have greatly advanced over the past decade and are transforming knowledge of neurotransmission in the nervous system^1–11^. Dopamine represents a key example: this neurochemical has received major attention due to its broad link to reinforcement learning, movement, and motivated behaviors^2,12,5^ as well as to neurological and neuropsychiatric disorders^13–17^. Dopamine is secreted within milliseconds following cell-membrane depolarization, is released into the extracellular space at synaptic or non-synaptic sites, and is transported across nanometer spaces in between cell boundaries to bind to receptors at neighboring cells^18^. Measuring these molecular dynamics precisely requires high spatial and temporal resolution. Physical sensors should occupy minimal dimensions to localize these molecular concentration changes in space as well as to reduce physical perturbation of the natural extracellular signaling environment. The size of the implanted sensor has been shown to correlate with the induced tissue response^19^, manifesting in the form of gliosis, scarring, and vascular infarction, which collectively reduce the sensor’s sensitivity to the surrounding neurochemical fluctuations^20^. Moreover, dopamine has been found to display spatially diverse signaling operations and functions, contrasting with past views of dopamine-mediated communication as globally homogeneous^5,21^. Measuring this spatial heterogeneity requires increases in the number of sensors, which increases the overall size or footprint of the implanted device. Methods to maintain a minimal implant footprint while simultaneously augmenting the total number of sensors for neurochemical monitoring could provide an important benefit.

Current methods to record neurochemical activity *in vivo* are usually in the form of genetically encoded fluorescent or electrochemical sensors. Genetically encoded fluorescent sensors, such as dLight and GRAB(DA), are recombinant proteins that fluoresce in response to binding of targeted molecules, such as dopamine^10,11^. These fluorescent sensors have been shown to display high molecular specificity and millisecond binding kinetics. Such sensors have enabled the detection of changes in dopamine concentration at high spatiotemporal resolution in a variety of organisms including flies, fish, and mice. The use of these fluorescent sensors requires genetic modification of the targeted neurons. Such genetic modification has shown limited success so far in primate species. Furthermore, fluorescent measurements in subcortical structures require insertion of optical fibers that are at minimum 100 µm in diameter and/or larger optical windows for 2-d imaging^1^, large enough to induce significant gliosis and breakage of blood vessels^3,4^.

Electrochemical methods, such as fast scan cyclic voltammetry (FSCV), avert the need for genetic modifications and large implant sizes associated with fluorescent sensors while providing the needed millisecond sampling and microscale spatial resolution^22,23^. FSCV as applied with implantable carbon fiber (CF) electrodes uses a low-amplitude voltage sweep (–0.4 – 1.3 V) to measure nanoampere-level current generated by voltage-dependent reduction and oxidation (redox) reactions of targeted molecules. The redox current is linearly proportional to the adsorbed analyte within a physiological range (nM – µM) allowing for quantitative predictions of *in vivo* neurochemical concentrations. CF materials have been established over several decades for FSCV-based recording of dopamine and other electroactive compounds. CF provides the optimal electrochemical interface due to its biocompatibility, as well as enhanced sensitivity, selectivity, and electron transfer properties^24^. Furthermore, these raw fibers are available off-the-shelf at diameters of 5 – 7 µm, which is essential to maintain the small dimensions of the final manufactured sensor. CF electrodes have been used to record millisecond dynamics of dopamine and other molecules in flies^9^, rodents^2–4^, monkeys^5–8^, and humans^25^. Critical limitations with this technique so far include the inability to distinguish similar compounds (e.g., dopamine and norepinephrine), and their restriction to mostly electroactive compounds (however, this can be addressed with enzyme coatings to convert non-electroactive compounds to electroactive H_2_O_2_^26^). Nevertheless, these techniques may present fewer safety barriers for clinical translation as they do not require genetic modifications to host cells, have been shown to operate over year-long time scales^3,5^, and have been used for recording in humans intra-operatively^25^.

A key advantage of FSCV-based neurochemical measurements is the ability to create ultrathin (∼10 µm diameter) and, as a result, highly flexible, implantable CF electrodes by leveraging the small diameters of the raw CF materials (5 – 7 µm) as well as conformal polymer coatings, such as parylene-C, for thin-film insulation^3,4,27^. The reduced diameter of these implanted sensors significantly reduces induced tissue responses and enhances functional longevity, enabling stable measurements of transient millisecond dopamine dynamics in rats for up to over a year^3^. Multi-channel approaches for neurochemical detection have so far been in the form of horizontal arrays containing individually penetrating CF electrodes that are laterally spaced with a pitch of 50 – 250 microns^4,27^. These horizontal configurations make brain insertion challenging for targeting deep subcortical brain regions due to the flexibility of these thin CF electrodes and the risk of individual electrodes buckling during brain penetration. Furthermore, the overall size of the array scales with the number of discretely spaced CF electrodes. The cranial opening must also be scaled in size to accommodate the array. Increasing the size of the cranial window increases the risks of surgical complications, trauma, and infection. Linear arrays that place sensors along the axis of insertion have been used widely for recording electrical neural activity in deep brain structures^28,29^. However, such vertical configurations limit sampling of neural activity to a single linear track in the brain. Methods that reduce the overall implant footprint (i.e., the size of the intracranial window) while providing access to transversely distributed sites would enable recording of the broader spatial dynamics of dopamine and other neurochemicals, as well as electrical spikes and local field potential neural signals.

Here, we report a novel CF electrode-thread (CFET) array that is inserted into the brain as a single tightly clustered bundle and splays with lowering to permit access to multiple laterally distributed subcortical brain sites. Clustering the electrodes into a tight bundle makes it possible to scale the number of channels while maintaining a minimal implant footprint. Our system leverages the innate flexibility of the thin parylene-coated CF (∼10 µm) electrodes to splay from their axis of insertion during cortical penetration and as lowered into deeper brain (5 – 20 mm). As a result, the sensors may be spread over a 2 – 18-fold larger area across subcortical brain targets compared to stiff electrode arrays. Furthermore, the electrodes are small enough to be sliced with the fixed brain tissue in post-terminal procedures, allowing precise localization of recorded sites relative to anatomical landmarks and any fluorescent or stained markers of protein expression. Assembly rigs were designed to enhance yield and throughput for fabricating the CF electrodes from the raw CF elements that make up the core neurochemical detection function of the device. Additionally, maltose was used to coat the CF electrodes in order to increase their rigidity during the implantation procedure. We constructed 8 – 16 channel CFET arrays and tested these *in vivo* in rats to validate multi-channel dopamine recording functionality, and with post-terminal histology to evaluate the spatial spread of the electrodes as well as to confirm that the positions of the CF tips remained intact during the thin-sectioning of the brain tissue.

## Results

### Design and Fabrication of CFET Arrays

CFET arrays were made up of multiple CFETs arranged in a tight bundle aimed to minimize overall implant footprint. The overall diameter of such a bundle is approximately equal to 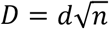, where *n* is the number of electrodes, and *d* is the diameter of the individual electrode. For example, a bundle of 10,000 electrodes of 10 µm diameter each would occupy an overall diameter of 1 mm. For comparison, the state-of-the-art Utah array (Blackrock) with 256 electrodes, versions of which are used widely for brain-computer interface (BCI) applications^30,31^, has a diameter of 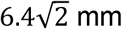 because of the discrete 400 µm spacing between individual electrode shanks. This inter-electrode spacing is designed to accommodate sampling of neural activity from a wider distribution of cortical sites at fixed intervals, which also necessitates a larger craniotomy and durotomy for implantation. By contrast, the compact CFET arrays introduced here were designed for targeting deep brain structures, where the thin and flexible properties of the individual CFETs could be leveraged so that they naturally separated and splayed during insertion. This splaying characteristic would then enable access to a wider range of sites from a smaller surgical cranial burr hole than the fixed arrays (**Fig. 1**).

**Fig. 1.**
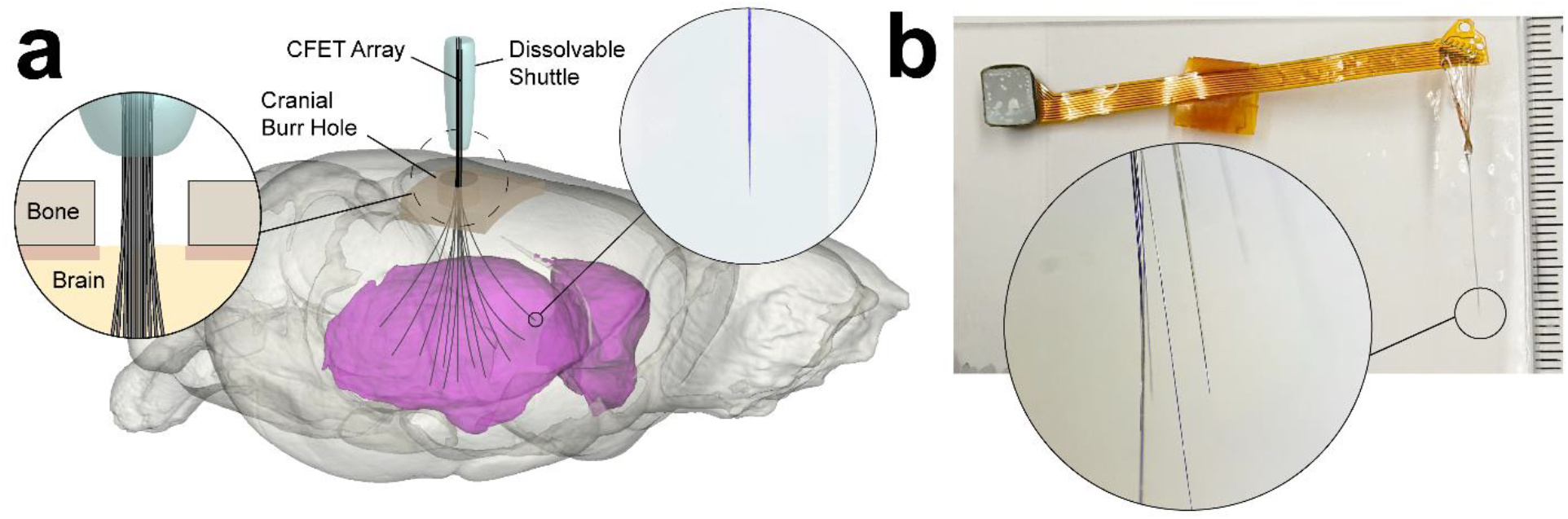
Illustration of carbon fiber (CF) electrode-thread (CFET) arrays. **a** CFETs are inserted into the brain (example shown is of a rat brain) through a single penetration point (i.e., cranial burr hole) as a bundle using dissolvable maltose as a temporary stiffener to increase insertion force at the tip and prevent buckling. Individual CFETs splay during the lowering process to spread to a lateral distribution of subcortical brain sites (e.g., within striatum, colored here in magenta). Each CFET has a discretely exposed CF tip for recording neurochemical activity. **b** Photograph of an assembled CFET array showing the bundled CFETs (right) connected to a flexible ribbon cable with a terminal press-fit connector for relaying signals to external FSCV instrumentation for recording neurochemical activity (left).

We fabricated CFET arrays of up to 16 patterned CFETs connected to a flexible printed circuit board (PCB) (flex-PCB) for interfacing with external FSCV recording instrumentation. CFETs and flex-PCBs were constructed separately. Fabrication of the CFETs involved the following steps (**Fig. 2**): Slotted aluminum fixtures were machined to tether bare CFs for subsequent parylene-C encapsulation (**Fig. 2a**). Double-sided polyimide tape was placed on both sides of the slot (25 – 35 mm) on the fixture that would be used to anchor the Cu wires and CF tips on either end. 10 – 20 Cu (40 µm diameter, annealed) wires (CU005200, Goodfellow Corp.) were cut to lengths of 10 – 15 mm each and were attached on one side of the slot, spaced 5 – 10 mm from each other on the pre-attached double-sided polyimide tape. Single 7-µm diameter CFs (C 005722, 34-700, Goodfellow Corp.), cut to 15 – 30 mm in length, were attached to the free-hanging ends of the Cu wires using silver epoxy (H20S, Epo-Tek), and the CF tips were then attached to the opposite end of the fixture slot on the pre-placed polyimide tape to anchor the CF tips.

**Fig. 2.**
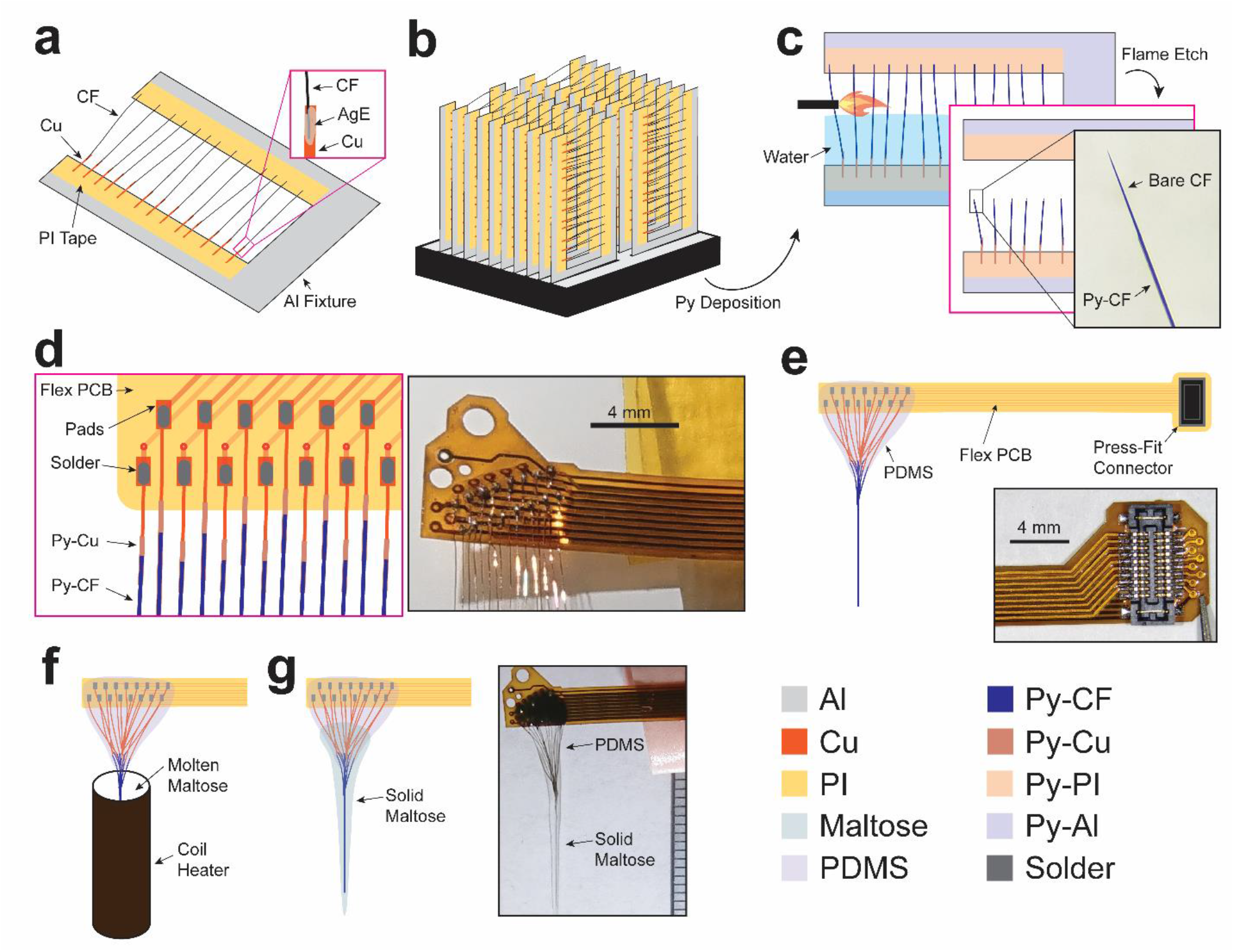
CFET array fabrication process. **a** Bare Cu wires are mounted onto an Al fixture using polyimide (PI) tape. The PI tape also serves to mask the tips of the Cu wires from subsequent parylene deposition in order that they can provide electrical contact for soldering steps. CFs are attached to the free-hanging end of the Cu wire with silver epoxy (AgE) and the opposite end of the CF is attached to the other side of the Al fixture’s slot with PI tape. **b** Al fixtures containing the Cu-CF assemblies are placed onto a standard microscope tray box, and then this tray is placed in a chamber for conformal vapor-deposition of parylene-C (Py) (1.5 – 3 µm thick). **c** The Py deposited Al fixture is immersed in water to thermally insulate the Py-CFs during flame etching to expose the bare CF tips (inset photograph shows a patterned Py-CF tip). **d** Etched Py-CF electrodes are tested *in vitro* and the bare Cu wire ends of the functional electrodes are soldered onto the flex-PCB shown as labeled illustration (left) and photograph (right). **e** PDMS is applied to the exposed pads of the flex-PCB and over the Cu wires to provide electrical insulation. In some devices, a thin layer of structural epoxy is applied onto the exposed pads before subsequent PDMS application. A press-fit connector is soldered onto the other end of the flex-PCB (inset photograph). **f** The completed CFET array is inserted into a temperature-controlled dip-coater containing molten maltose and the array is lifted from the maltose at a controlled rate to provide a 1–3 mm thick coating of solid maltose. **g** Final maltose-coated CFET array (inset shows photograph of the same) ready for *in vivo* use.

These long CFs are extremely flexible (1/*k* = 4*L*^3^/(3*πEr*^4^), where *k* is stiffness, *E* is the Young’s modulus of the CF ∼ 200 GPa, *r* is the radius, and *L* is the length) given the ultra-high aspect ratios (length to width ratio of > 2,000) applied here to reach deep subcortical brain targets. A CF will deflect and even attach to nearby surfaces due to air currents caused by its manual repositioning, movements in the room, and room ventilation, as well as electrostatic forces. The purpose of the fixture’s slot was to keep the samples anchored, yet elevated and separated to insulate, conformally, the individual Cu-CF assemblies with parylene in subsequent vapor-deposition steps. Standard microscope slides were also used successfully as fixtures where the CF tips were left unanchored and remained elevated vertically above the slide. However, this required greater spacing between Cu-CF assemblies in order to prevent neighboring CFs from contacting and adhering to each other, which would ultimately limit the total number of Cu-CF assemblies that could be placed in the chamber for parylene deposition. Each sample-loaded fixture was placed in an oven to cure the silver epoxy at 80 – 90 °C for 2 – 3 hours. The fixtures had similar dimensions to a standard microscope slide (1/16 inch thick) and were stacked in a standard microscope container tray (82024-606, VWR) that had been trimmed to fit in the parylene deposition chamber (PDS2010, Specialty Coating Systems) (**Fig. 2b**). All Cu-CF assemblies were treated in an adhesion promoter solution consisting of 3-(Trimethoxysilyl)propyl methacrylate (Silane A174, 440159, Sigma-Aldrich), isopropyl alcohol, and distilled water at a volumetric ratio of 1:1000:1000. Parylene was deposited onto the fixtures containing the Cu-CF assemblies to a thickness of 1.5 – 3 µm (Fig. 2C). Parylene-coated CFs (Py-CF) were flame etched using a butane torch to expose a discrete length of CF at the tip, following methods established in previous work^3–5,32^ (**Fig. 2c**). The level of water relative to the base of the CF controlled the final lengths of the individual electrodes that would be inserted into the brain. The lengths of etched Py-CFs ranged from 5 – 22 mm to reach subcortical brain structures (e.g., striatum) in rats and to target emulated depths of the primate striatum (> 15 mm) in agar brain phantom experiments.

Etched Py-CF electrodes were then tested *in vitro* in saline to record the background current using FSCV and measure targeted functional metrics for sensitivity (i.e., maximum background current) and limits of detection (e.g., noise) to dopamine^3^. Targeted metrics were 500 – 800 nA for background current, and < 0.05 nA for noise, as these were empirically determined to provide optimal performance in past *in vivo* measurements^3–5,8^. Cu-CF electrodes meeting these specifications were aligned on a slim-profile custom-made flex-PCB so that the bare-ends of the Cu wires lay on the exposed conductive pads and were temporarily held in place with double-sided polyimide tape (**Fig. 2d**). Solder paste (SMD291AX, Sn63/Pb37, Chipquik) was applied to the pads and Cu wires, and then a hot-air gun was used to melt the solder and create conductive bonds. The other end of the flex-PCB was terminated with a press-fit connector (Slimstack 5024262030, Molex) to plug into our custom-made FSCV instrumentation and relay signals to and from the CF electrodes for neurochemical recording^4^. Polydimethylsiloxane (PDMS) (Sylcap 284-F, Microlubrol) was applied over the exposed conductive pads and Cu wires to insulate these traces (**Fig. 2e**).

We also attempted an alternative method to construct the CFET arrays, aiming for faster throughput. This method involved soldering all 16 Cu-CF assemblies to the flex-PCB first, along with the press-fit connector. The resulting flex-PCB with attached Cu-CFs was then secured to an aluminum fixture, with the flex-PCB mounted on one end and CF tips on the other, using double-sided polyimide tape. The entire array was subsequently processed through parylene deposition and flame-etching. However, *in vitro* testing displayed a lower yield of functional channels with the completed array due to incomplete etching of some of the Py-CF tips. This incomplete etching occurs because air flow from the torch pushes the exposed Py-CF tip into the water before it can be exposed to high enough temperatures from the center of the flame for sublimating the parylene. The initially described fabrication procedure produced a significantly greater yield (> 90%), as only functional Py-CFs were selected to solder onto the ribbon cable.

Dissolvable maltose was coated onto the CFET array to stiffen the bundle to facilitate subsequent brain insertion (**Fig. 2f**). A custom-made temperature-controlled dip-coater was used to coat the electrodes reproducibly with maltose. Nichrome (80% Ni, 19.5% Cr, 1.45% Si) wire (20G, Master Wire Supply) was coiled around a hollow ceramic tube. A thermistor was attached to the base of the tube to monitor the temperature of maltose inserted inside the tube, and then the base was sealed to prevent maltose from leaking using thermally-conductive epoxy (832TC, MG Chemicals). A solid-state relay was used to provide DC current (up to 1A) to the heating wire. An Arduino was programmed to switch the current on and off based on comparing the temperature measured from the thermistor to the set-point (110 – 120 °C). Maltose powder (M5885-100G, Sigma-Aldrich) was placed into plastic falcon tube vials and then submitted for gamma-irradiation sterilization (25 kGy, VPT Radiation Lab and Test Services).

The CFET array and coiled heater were sterilized using vaporized hydrogen peroxide (Sterrad). The array was lowered into the tube and then maltose powder was poured into the heating tube, and heated until molten. The temperature was lowered by 2 – 5 °C to increase the viscosity and after a few minutes the array was lifted at a rate of 250 µm/s. This procedure produced a relatively uniform thickness of 1 – 2 mm around the bundle, but success was highly dependent on the volume of maltose inside the tube due to the non-uniform temperature distribution across the coil. Alternatively, we also employed a manual coating method for the CFET array. We first melted maltose powder on a hot plate over which CFET array was placed and rapidly lifted to produce a relatively uniform coating of 1 – 3 mm diameter.

### Recording Sub-second Molecular Dynamics of Dopamine *In Vivo*

*In vivo* functional neurochemical recording operation of the arrays was tested in an anesthetized rat (**Fig. 3**). Drilling a small burr hole (1.5 mm diameter) in the skull was sufficient to allow channeling of the arrays to multiple (up to 16) spatially distributed subcortical sites in the striatum. The array was inserted into the brain using a dissolvable maltose shuttle, which rigidly fixes the CFETs just above the exposed cortical surface to generate sufficient penetration force. The array was usually not lowered until ∼ 1 mm of the rigid maltose shuttle above the brain surface had been dissolved by applying drops of saline. The array was inserted to a depth of 6.5 mm DV targeting the ventral striatum.

**Fig. 3.**
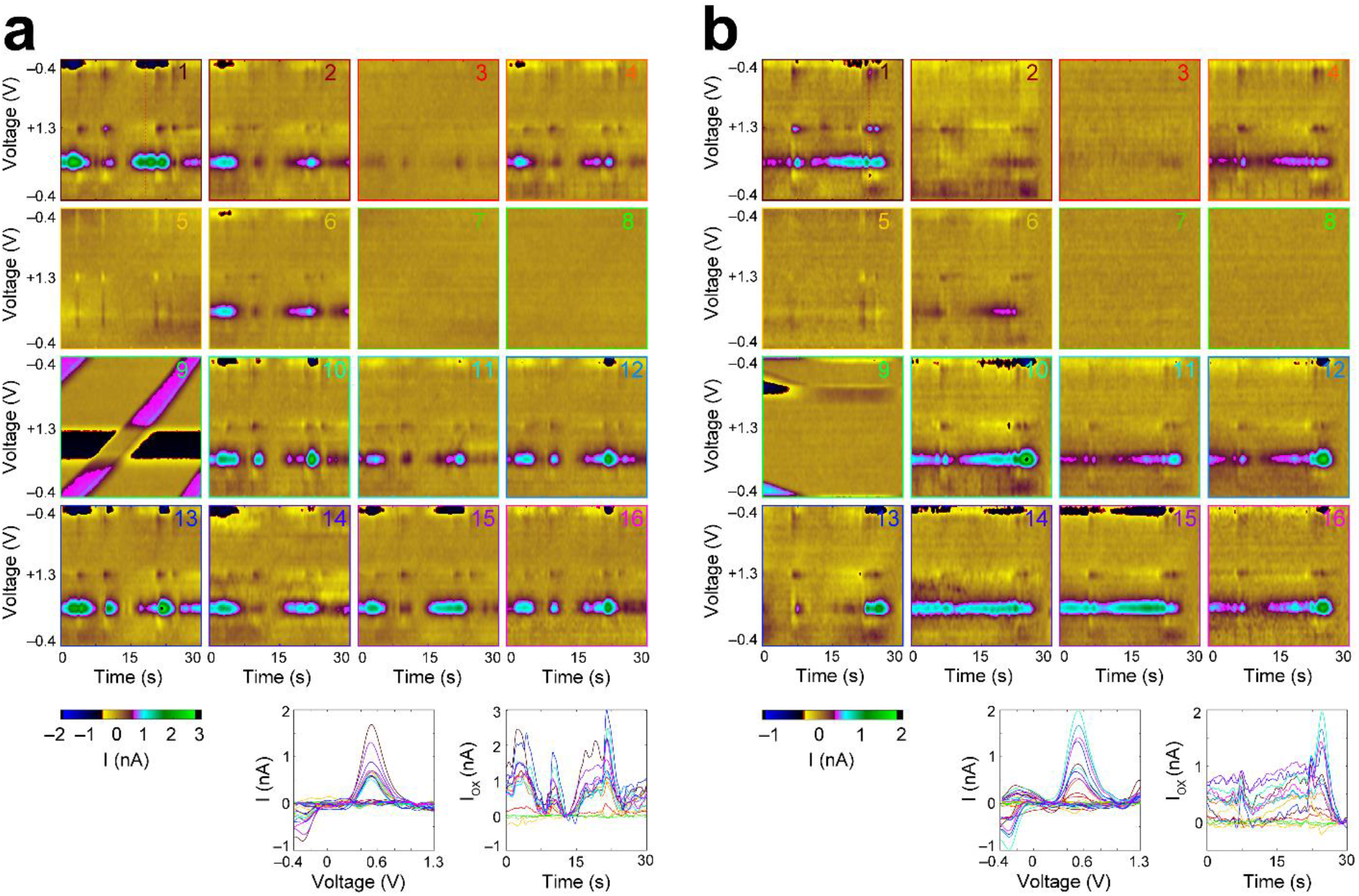
*In vivo* measurements of pharmacologically induced dopamine transient signals from the implanted CFETs in the rat striatum. **a** (Top) Background-subtracted color plots (x-axis is time, y-axis is voltage applied, color = current) of FSCV recording showing current fluctuations along dopamine redox (measured at –0.3 and 0.55 V) potentials, 33 minutes following administration of raclopride and cocaine. Multiple channels (1, 2, 3, 4, 6, 10, 11, 12, 13, 14, 15, 16) display significant dopamine concentration changes (channel numbers indicated on top right of each color plot). Channel 9 is a reference channel for noise removal and displays the line noise that was uniformly subtracted from all other channels (Bottom-Left) Cyclic voltammogram (CV) showing peak current changes along dopamine redox potentials to illustrate molecular identity of measurements. (Bottom-Right) Dopamine oxidation current (i.e., current at oxidation potential ∼ 0.55 V), which directly correlates with dopamine concentration, as a function of time. **b** Same as (**a**) for a time window 139 minutes following drug administration displaying greater dopamine signaling heterogeneity across the channels.

We recorded pharmacologically-evoked dopamine transient signaling in the striatum using FSCV in most (12/15) of the implanted CFETs (one of the channels was disconnected due to poor pin connection at headstage). Endogenous dopamine transients are rarely detected in the striatum of the anesthetized rat. Therefore, most studies use electrical stimulation and/or dopamine-selective drug treatments to evoke dopamine signaling^4,8,33^. We administered a combination of raclopride (3 mg/kg) and cocaine (15 mg/kg)^33^, respectively a dopamine D2-receptor antagonist and a dopamine transporter inhibitor, which act to increase dopamine transient signaling in the striatum^34^. Dopamine transients were observed within 5 minutes after I.P. injection.

The transient dynamics observed in our measurements were similar to those reported in previous studies using the same drug combinations^34^. All of the recorded implanted CFETs (14/14) displayed optimal functional characteristics in terms of noise (i.e., < 0.05 nA), which defines the absolute limit of detection. Four of the CFETs exhibited lower background currents than the level (500 nA) we have previously established empirically to provide successful dopamine measurements (channels 3, 6, 7, 8), but two of these CFETs (channels 3 and 6) still were able to detect visible dopamine transients in the form of selective changes in dopamine redox current (**Fig. 3a** and **b**). One of the CFETs had been cut bluntly at the tip with scissors to minimize its conductive surface area in order to use it as a reference channel for subtracting external electromagnetic interference (EMI) line noise common to all electrodes. This reference electrode had a current ∼ 64 nA and noise < 0.01 nA. These results confirm functional operation of the arrays in detecting millisecond neurochemical dynamics in multiple sites of the striatum and as implanted through a small (1.5 mm) cranial burr hole. Both raclopride and cocaine have been demonstrated to increase burst firing of dopaminergic neurons and to generate increases in the resulting frequency and amplitude of transient dopamine release events in the striatum^7^. In this study, we were able to observe such events using our implanted CFET arrays.

The surgical implantation procedure was considerably more straightforward compared to previously reported multi-channel neurochemical arrays^3,4,27^. Previously reported arrays were horizontally configured, meaning that individual electrodes were inserted through separate penetrations and required a wider craniotomy to accommodate multiple electrodes spaced along the transverse plane. Ideally, all of the electrodes should be inserted normal to the brain surface and vertically with respect to the axis of insertion to ensure maximal penetration force and reduce the chance of buckling. However, the flexible electrodes will curve and bend slightly during the process of coating them with a dissolvable shuttle (e.g., PEG or maltose), making it difficult to maintain a vertical entry angle for the electrodes. It is usually difficult and/or impossible to find a position over the brain where all of the electrodes have clear access to the overlying brain parenchyma away from superficial blood vessels. These larger blood vessels are significantly stiffer than the surrounding brain tissue and impenetrable by thin CF electrodes. In some instances, electrodes must be manually guided and/or straightened during the insertion process, which demands skilled dexterity. These challenges scale with the number of electrodes that must be manually guided into the brain. A tightly clustered array, as demonstrated in this work, allows for a single insertion, reducing the number of individual entry points and penetrations into the brain and consequently the overall likelihood of failure due to buckling. This improvement is especially noteworthy given the greater depth of our targets in comparison to most other cortical applications of reported CF arrays^27,32,35–37^ and/or use of guide cannulas to shuttle the probes to deeper brain areas^27^. However, we should note that our maltose coating procedure does not always provide a uniform thickness radially around the diameter of the CFET array. A non-uniform coating may cause electrodes on one edge to be released from the shuttle faster than adjacent perimeters of the array. Dissolving only the basal (∼ 1 mm length) component of the coated array while keeping the rest of the coating intact requires a responsive control over the insertion speed.

Our array fabrication procedure is highly scalable, allowing for a substantial increase in the number of channels for capturing dopamine’s spatially diverse operations^5,21^. Specifically, our batch etching protocol and customizable slim-profile flex-PCBs were made to be high-throughput and thus readily allow expansion to accommodate more channels.

### Characterization of Spatial Spread of Arrays as a Function of Insertion Depth *In Vitro*

Individual CFETs were inserted into agar phantoms emulating the mechanical properties of the brain to characterize deflection as a function of insertion depth and tip profile (i.e., flame-etched and blunt-cut) (**Fig. 4a**). CFETs were successfully inserted to depths of greater than 20 mm to visualize trajectories that would be used to reach deep brain structures in rodent and monkey brains (e.g., the dorsal edge of the striatum is 13–15 mm from the brain surface in the Rhesus monkey). The maximum deflection observed at a depth of 5 mm, equivalent to reaching the middle of the striatum in rats, was 0.81 mm for the flame-etched and 2.12 mm for blunt-cut electrodes. The maximum deflection observed at a depth of 18.5 mm, equivalent to placements within mid-depth of the striatum in the Rhesus monkey, was 2.1 mm for the flame-etched and 5.5 mm for the blunt-cut electrodes. The spatial deflections observed for both CFET types would allow areal coverage up to 4.6 mm^2^ (1.72 mm^2^ for flame-etched) in the striatum in rats and 137.17 mm^2^ (15.72 mm^2^ for flame-etched) in monkeys at the aforementioned depths as estimated from linear fit (see **Fig. 4b** and **c**). These results predict an estimated 2.2 – 17.5-fold increase in areal coverage relative to stiff electrodes that may be accommodated within a 1 mm diameter burr hole (0.78 mm^2^ areal coverage).

**Fig. 4.**
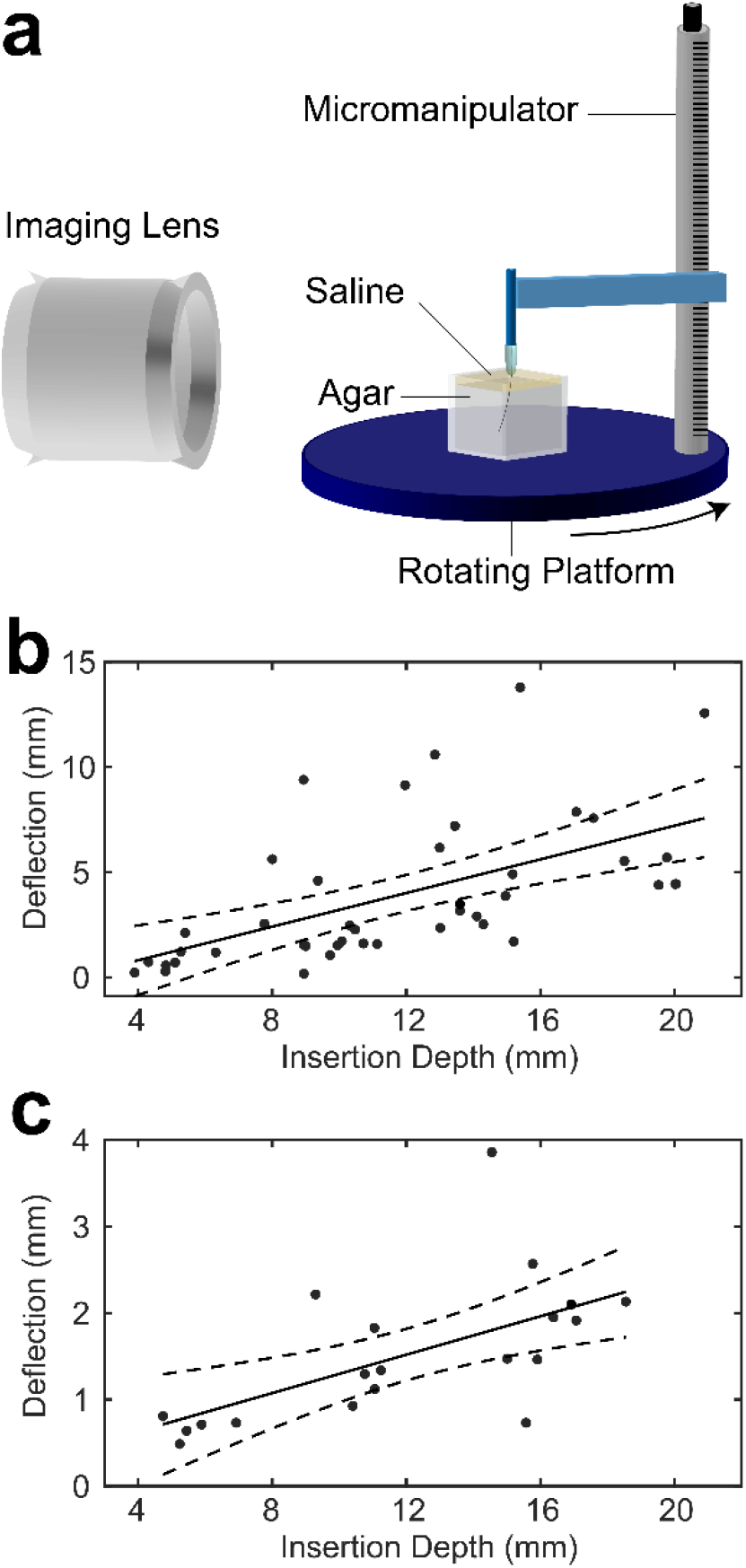
*In vitro* CFET splay characteristics. **a** Apparatus for measuring deflection in agar brain phantom. A CFET is anchored to the micromanipulator arm and is contained within a maltose shuttle that dissolves as lowered into the saline, replicating the *in vivo* insertion procedure. **b** Deflection vs. insertion depth for blunt-cut CFETs. The solid line is the linear fit with a beta coefficient of 0.40 and an intercept of –0.79 mm (R^2^ = 0.326, p = 6.2e–05) and the dashed lines represent the 95% confidence bounds for the regression slope and intercept. **c** Same as (**b**) for flame-etched CFETs with a beta coefficient of 0.11 and an intercept of 0.18 mm (R^2^ = 0.385, p = 0.002).

The magnitude of deflection was observed to be significantly less with flame-etched tips (deflection/insertion ratio of 0.128) than with manually-cut tips (deflection/insertion ratio of 0.312) in the agar brain phantom. Flame-etched tips display axisymmetric conical tips that experience theoretically uniform cutting forces along the transverse plane. Deflection may arise due to a combination of factors including the angle of initial penetration into the gel phantom as the maltose coating may have generated subtle bends in the electrode shaft when it was applied. Another possibility is the asymmetry of frictional forces along the electrode shaft during insertion^38^. Manually-cut electrodes display high bevel tip angles (80 – 90 degrees). Lateral bending forces increase with bevel tip angle as described by the following equation, F_L_ = F_Z_ tan(θ), where F_Z_ is the axial force and θ is the tip angle relative to insertion axis. Empirically, the higher degree of deflection observed for blunt-end tips accords with the deflection characteristics measured for small 60 µm capillaries inserted into brain phantom and tissue^39^. Flame-etched electrodes are in many cases further trimmed with a razor to reduce the exposed CF surface area to bring the background current to the targeted range of 500 – 800 nA. Such trimming increases the angle of the tip to that of the blunt-cut profile, and therefore deflection would also likely increase. Insertion in the agar brain phantom does not replicate the inhomogeneous mechanical properties of the real brain parenchyma such as the different characteristic elastic moduli of gray and white matter^40^, the interspersed network of blood vessels, and the existence of a vast array of molecules and proteins that would affect the relative viscosity of the surrounding fluid as well as the friction experienced by the electrodes during tissue insertion^41^. The tight configuration of the electrodes with interleaving fluids (i.e., dissolved maltose, exogenously applied saline, and brain cerebrospinal and interstitial fluid) would also induce capillary forces between the electrodes that would act to hold them together and reduce relative splay. Therefore, future *in vivo* or *ex vivo* studies are necessary to more accurately estimate the deflection characteristics given the aforementioned variables that are not accounted for in our *in vitro* brain phantom.

### CFETs Fixation and Cutting of Sections within Brain for Precise Localization of Recording Sites

Locating the implanted electrodes in the brain tissue is critical to identify relevant anatomical and molecular markers that characterize the recorded site using post-terminal processes such as standard immunohistochemistry. Yet, most methods to locate the recording sites have limitations in spatial resolution (> 100 µm) and/or difficulties in distinguishing them because of their small size and/or resemblance to natural anatomical structures such as blood vessels in the perfused brain. Larger electrodes (> 100 µm) displace and/or cut the tissue during insertion creating a noticeable “track” or hole similar in diameter to the electrode itself. Further, these larger electrodes produce significant tissue responses in the form of neuroinflammatory responses, gliosis, and perforation of blood vessels–processes that can extend hundreds of microns from the recording site. Such processes are usually observed through heightened expression of astrocytes, microglia, and blood brain permeability (i.e., IgG) using standard immunohistochemistry. These processes obscure spatial accuracy of the recording site due to the size of the induced mark (> 100 µm) as well as the ability to evaluate unperturbed anatomical and molecular markers in its vicinity. Electrical lesions are also used by applying sustained DC current (> 10–15 µA) at the electrode tip *in vivo* to burn and mark the tissue, which also destroys the natural environment surrounding the recording site and increases tissue responses^3^. On the other hand, smaller electrodes, such as variations of CFET electrodes including our own (< 15 µm)^3,4,27,32,35,36^ do not produce visible inflammation after chronic implantation unless electrolytic lesioning is used and generate “tracks” that when visible are largely indistinguishable from blood vessels in the perfused brain.

We developed electrode-tissue retention methods to allow direct reconstruction of the location of recording electrodes in the brain tissue by leveraging the ultra-thin characteristics of our electrodes. An advantage of the CFET is its small diameter (∼10 µm), which necessarily reduces shear strength and the needed cutting force that would allow for cutting properties similar to that of the surrounding brain tissue. We used a standard microtome to directly cut 100 µm-mm thick sections with the electrodes still embedded in the fixed brain tissue. This method allowed retention of their exact placement in the brain. We confirmed that the position of electrodes retained in the sliced brain tissue coordinates with the location of focally enhanced IgG expression that are typically used to identify electrode tips in acute implantation procedures. Blood vessels at the brain surface were deliberately penetrated in order to convey markers of blood-brain barrier leakage (i.e., IgG) with the brain inserted electrodes. Slicing of the integrated CFET-tissue volume further allowed direct reconstruction of implanted CFET locations relative to neurochemical landmarks in the striatum (e.g., striosome vs. matrix in the dorsal striatum) (**Fig. 5**).

**Fig. 5.**
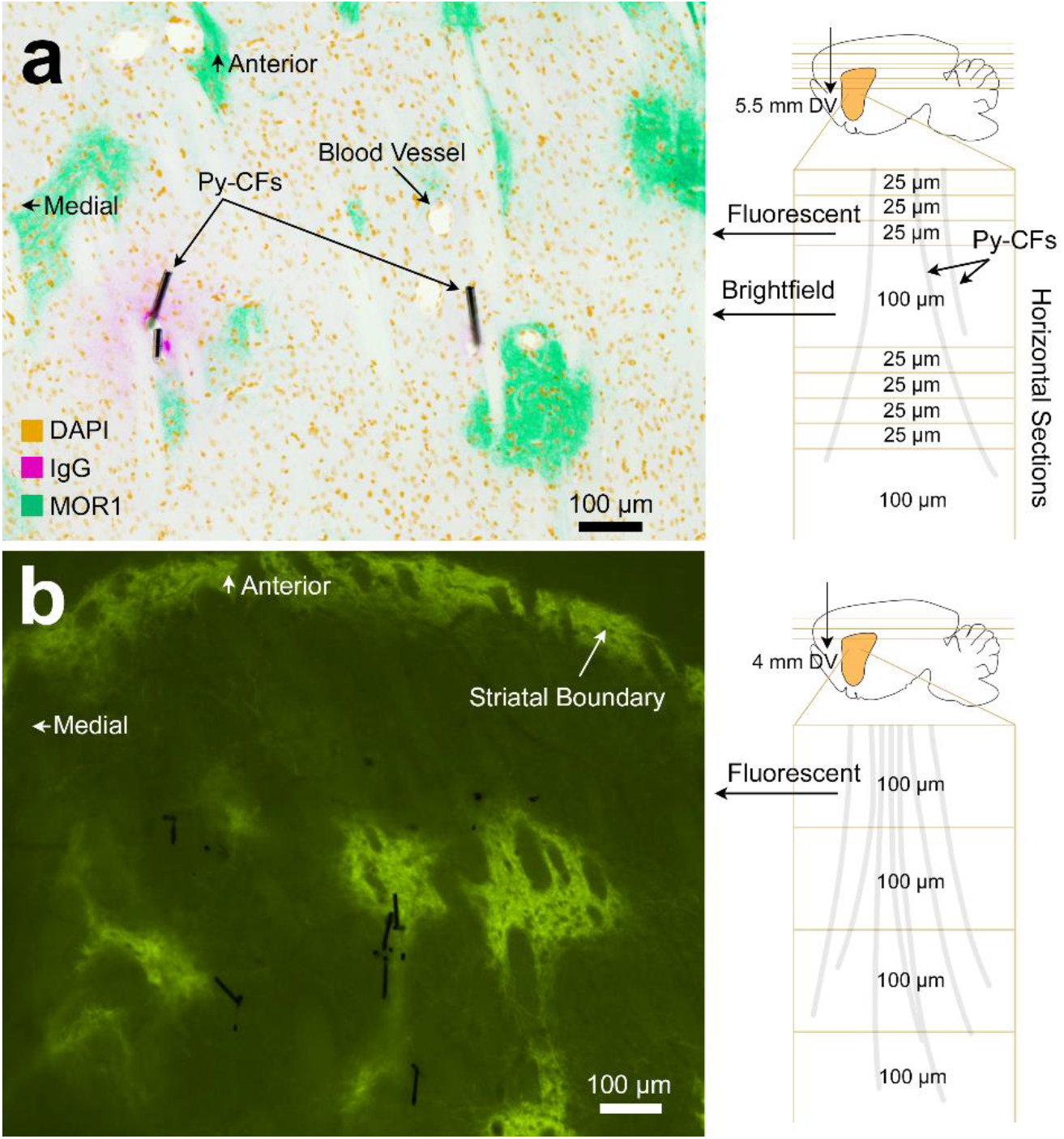
Tissue retention of CFET arrays. **a** (Left) Bright-field image of a horizontal section through the brain of a rat, illustrating two examples of Py-CFs maintained in the brain, overlaid with a reverse-contrast fluorescent image of the section just above that was stained for markers (color key in figure) of cell nuclei (DAPI), blood brain permeability marker (IgG), and striosomes (MOR1). The Py-CFs were unable to remain in the thinner 25 µm slices typically used for immunohistochemical staining, and therefore sections were made thicker (100 µm) in some sections (sectioning schedule shown in a, right). Enhanced IgG staining is apparent just around the Py-CFs, confirming that the devices were not displaced during brain cutting. Blood vessels were perforated at the surface of the brain during insertion of the CFET array in order to provide staining along the CFET insertion track and to validate that the location of the Py-CFs remained fixed in the tissue section. **b** Fluorescence image of a MOR1-immunostained 100 µm thick section with embedded Py-CFs (black lines).

Similar methods were developed recently to retain similar CFET devices in brain tissue slices^27^. These previously reported methods, however, required use of a decalcifying agent to dissolve the skull. It was not clear whether the electrodes may have moved between processing steps, as no validation was done to ensure that their placement remained secured, and their staining procedures were only done on adjacent tissue slices from those embedded with the CFETs. Our procedure as outlined here is relatively simple; the device was simply cut through with a rotary saw, after which routine removal of bone using rongeurs or other bone clipping instruments could be performed, followed by standard histology protocols to slice and stain the brain tissue. Furthermore, we were able to stain the same tissue section that contained the CFETs with markers indicating expression of MOR1, used to differentiate striosome and matrix sub-compartments in the striatum. Imaging these molecular markers on the same slice as the CFETs would enable maximal accuracy with respect to characterizing the neurochemical and anatomical profiles of the cells directly adjacent to and defining the recording site. Tissue sections were made relatively thick (100 µm) to ensure that CFETs remained in the tissue during multiple shaking, washing, and staining steps used to prepare sections for immunohistochemistry.

A remaining challenge is functionally mapping the retained electrode tips with the associated functional measurements given their random spatial distribution in the CFET array. One possibility to address this may be to apply fluorescent dyes and/or stains of different emission wavelengths to each electrode to discern their channel configuration from the functional measurements^42^.

## Discussion

We demonstrate a novel splaying array for navigating to multiple subcortical sites that are laterally distributed from the insertion axis to record neurochemical signaling dynamics. Key advancements include the ability to access widespread neural targets while eliminating the need for a large cranial window. Dopamine measurements in humans to date have only been performed successfully in acute intra-operative contexts^25^. Minimizing the cranial window and the size of excised dura mater reduces risks of trauma and infection, facilitating translation of neurochemical sensors to clinical applications for diagnosis and closed-loop treatment (e.g., deep brain stimulation) for a wide range of disorders, including Parkinson’s disease. Recent technological advancements in multi-site recording of dopamine have shown that dopamine and other neuromodulator molecules operate much more heterogeneously than previously thought^5,8,10,11,21,43^. A strong motive for the technology presented here was to develop new tools that would allow spatial mapping of these dynamic neurochemical signals to characterize the function of their heterogeneity. Another key capability of the CFET array, not evaluated in this work, will be to enable stable dopamine neurochemical recording over extended time periods that encompass more gradual and adaptive processes, such as multi-stage skill learning. Our CFETs manifest similar characteristics to the micro-invasive electrodes shown to preserve stable recording functionality for over a year in rodents and monkeys^3,5^. Nevertheless, chronic implantation and testing to ensure the integrity of hermetic encapsulations and insulation of electrical components will be necessary to validate the same claims for the device presented here. CF electrodes, such as those used in this work, are capable of recording electrical neural activity^27,32,35–37^–functions not directly tested in this study. The dual functional capabilities of these CF-based sensors for both electrical and chemical recording merit a critical advantage over most standard materials used for electrophysiological measurement (e.g., platinum-iridium, tungsten, and gold). Development of such multi-modal sensors is essential to allow discovery of the effects of neurochemical fluctuations on neuronal spiking activity that ultimately regulates behavior and to uncover functional mechanisms allowing neuroplasticity over time.

## Methods

### FSCV Methods for Neurochemical Recording of Dopamine

An integrated system of hardware and software capable of recording dopamine signals up to 16 channels simultaneously using fast scan cyclic voltammetry (FSCV) was built with off-the-shelf electronics and a PC based data acquisition system^4^. Guidance on constructing this system, including the circuit board layout as well as the MATLAB software for visualizing dopamine recording in real-time can be found on the GitHub Repository (https://github.com/hschwerdt/multifscv).

FSCV is an electrochemical method widely used with CF electrodes to measure chemicals in the brain with high temporal (i.e., milliseconds) and spatial (i.e., micrometers) resolution. The optimal waveform for detecting dopamine and other catecholamines has been empirically established and involves applying a triangular waveform ramping from –0.4 V to 1.3 V at a scan rate of 400 V/s^44^. These waveforms are applied at a rate of 10 Hz and held at –0.4 V between scans. Dopamine molecules near the CF undergo oxidation (converted to dopamine-o-quinone by losing electrons) and reduction (converted back to dopamine by accepting electrons). Such oxidation and reduction (i.e., dopamine redox) process leads to current changes at –0.2 V and 0.6 V relative to an Ag/AgCl reference electrode.

The CF electrodes are conditioned before recording by applying the same ramping voltage at a cycle frequency of 60 Hz for 15 minutes in order to stabilize changes in background current. The recorded current change along the applied voltages at each scan is concatenated in order to create background-subtracted color plots (**Fig. 3**). These plots use a nonlinear color scale to represent current, with the x-axis indicating time and the y-axis indicating applied voltage. At each scan, the cyclic voltammogram (CV) is computed to identify the redox potentials of dopamine (i.e., –0.2 V and 0.6 V). These redox potentials may shift due to dechlorination of the Ag/AgCl reference, *in vivo*. Therefore, an offset potential is applied from 0.1 to 0.2 V in an attempt to shift observed dopamine redox potentials. All of these analyses were performed in MATLAB (MathWorks, MATLAB, 2020) as previously reported^3,4^.

### *In Vitro* FSCV Testing to Characterize Functional Operation in Dopamine Detection

Each etched CF electrode was tested *in vitro* in a beaker containing 0.9% sodium chloride (saline) to determine its functional properties (i.e., background current and noise) before soldering to the flex-PCB. *In vitro* testing was performed in a Faraday cage to minimize EMI interference during test recording. A custom designed PCB adapter was used to temporarily attach and connect up to 16 etched Py-CF assemblies for FSCV testing, in some tests. Py-CF threads were determined to be suitable for *in vivo* recording if they met both of the following 2 criteria: (1) current noise < 0.05 nA, and (2) magnitude of background current was in the range of 500 – 800 nA, which corresponds, respectively, to the limit of detection and the sensitivity to dopamine, as established previously^3^. A steel razor blade was used to trim the tip of the exposed CF slightly in instances in which the magnitude of background current was above 800 nA, and then the electrode was retested. The trimming and test process was repeated until the final background current was in the target range of 500 – 800 nA. The Py-CF electrodes were determined to be nonfunctional and would not be trimmed if they met any of the following 2 criteria: (1) a mechanical break of the electrode along its shaft or etched tip that resulted in a measurement of background current less than 100 nA, or (2) a perforation of the electrode that resulted in current noise ≥ 0.05 nA or background current saturation (magnitude of background current at any potential ≥ 2000 nA). This protocol can also be found at protocols.io (dx.doi.org/10.17504/protocols.io.rm7vzbdxrvx1/v1).

### *In Vivo* Device Implantation and Testing Procedures

All procedures involving animals were approved by the Committee on Animal Care at the Massachusetts Institute of Technology and were performed strictly following the U.S. National Research Council Guide for the Care and Use of Laboratory Animals. Long–Evans male rats were used for tissue retainment testing (n = 2) and dopamine recording (n = 1).

Rats were first anesthetized (1.5–2.0% isoflurane, 1 L/min oxygen) and were administered an analgesic (Meloxicam, 2 mg/kg) subcutaneously. Rats were fixed in stereotactic frames (Stoelting, 51600, 51449) that carried vertical micromanipulators used to navigate to targeted brain areas. A sterile field was created on the skin above the cranium. The skin was incised and retracted to expose the cranium. The CFET array was targeted to be placed in the right striatum (anteroposterior [AP] +0.5 mm, mediolateral [ML] +3 mm). The reference Ag/AgCl electrode, made of an insulated silver wire (A-M Systems, 787000 and 786000) with an exposed 0.5–1 mm tip chlorinated in bleach overnight, was targeted to be placed in the contralateral hemisphere, in the cerebellum. In addition, 5 – 10 bone screws (Stoelting, 51457) were placed on the circumference of the exposed calvarium in order to provide additional connections for electrical grounding as well as to secure subsequent cement (for the tissue retainment tests). A bare stainless-steel wire (A-M Systems, 792900) was wound around 2 or 3 screws to serve as an electronic ground connection.

The dura mater overlying the site calculated to be at the A-P level of striatum was removed with the bent tip of a 32G hypodermic stainless-steel needle. Afterwards, the micromanipulator was manually driven at a rate of 0.1 – 0.3 mm/s to lower the maltose coated CFET tips from the starting position. After the maltose coated CFET tips approached just above the brain surface, a small amount of saline was applied to dissolve ∼ 1 mm length of the basal portion of the maltose coating of the CFET tips. The non-maltose-coated portion of the CFETs was then lowered into the brain. The penetration of CF into the brain was supported by the rigid maltose coating above the basal portion. Afterwards, the process of saline dissolving and CFET array lowering was incrementally repeated until the array was lowered to a final depth of 4 – 6.5 mm dorsoventral (DV) relative to the cortical surface. The implanted array was connected to the FSCV headstage for dopamine recording experiments. For brain fixation and CFET sectioning experiments, the implant was secured to the skull by applying acrylic cement (Ortho-Jet, 0206) over the array and the skull. This was followed by protocols described below, in “<u>Histology and Retention of Implanted Electrodes in Fixed Tissue”</u>. FSCV recording began once the array reached its targeted depth and continued for a period of 15 – 30 minutes until the background current had stabilized (i.e., < 1 nA change in a 30 s window for most of the functional channels). 3 mg/kg raclopride and 15 mg/kg cocaine were administered to the anesthetized rat intraperitoneally to induce dopamine transient signaling in the striatum. Dopamine transients were observed five minutes after the injection of raclopride and cocaine. Recording continued for 2 hours after the drug administration. Chemical selectivity to dopamine was confirmed by the presence of sharp current peaks at the known redox potentials for dopamine (∼ –0.2 and 0.6 V) as well as by the selective action of the administered drugs acting at dopamine receptors and transporters. The inserted depths were DV 6.5 mm for the functional recording experiments (**Fig. 3**), and 5.5 mm (**Fig. 5a**) and 4 mm DV (**Fig. 5b**) for the tissue retainment studies. These methods are also described at protocols.io (dx.doi.org/10.17504/protocols.io.14egn2q3qg5d/v1).

### Characterization of Splay Characteristics in Agar Brain Phantom

The spatial insertion profiles of the electrodes were emulated by inserting individual electrodes into transparent agar phantoms where electrode trajectories could be visualized. We characterized electrodes with two different types of tip profiles: flame-etched tips displaying a conical CF end profile, and razor-cut tips displaying a large tip angle usually close to 90 degrees but this could vary depending on the alignment of the cut. These are the two main profiles that have been applied in prior work for neurochemical and electrical neural recording^3,4,27,32,35^ and are expected to represent the two extremes in terms of extent of generated deflection angles^39^.

Agar phantoms were made using 0.6% agar (Sigma-Aldrich, 05039-500g) in distilled water to emulate the mechanical properties of brain. Agar was dissolved in boiling distilled water (100°C), covered, and allowed to mix for 10 minutes at 80°C, using a magnetic stirrer and hotplate, until the solution looked visibly homogenous. The solution was cooled at room temperature for half an hour, before being transferred into transparent plastic cells (open cubes with side lengths of 4.5 cm) for penetration testing. Agar containing cells were refrigerated overnight at 20°C to solidify. Agar phantoms sat at room temperature for a few hours until the phantom reached and were maintained at room temperature before testing (IR5 Dual Laser Infrared Thermometer, Klein Tools).

The setup for deflection characterization consisted of a micromanipulator for holding and lowering the individual CFET to be tested, a rotating plate that held the agar phantom and manipulator and allowed viewing from all angles, and a stereomicroscope with a camera to view the trajectory of insertion and measure the deflection distance (**Fig. 4a**). All tests were performed with maltose-coated CFETs. Agar phantoms were covered with a thin layer of saline to dissolve the maltose as the electrode penetrated the agar phantom, and the saline was dyed yellow with food coloring to help distinguish the agar-saline interface and to identify when penetration had been achieved.

A maltose coated electrode was attached to the micromanipulator and lowered manually into the saline-covered agar phantom. Manual lowering was used to ensure that the maltose was properly dissolved at the saline/agar interface before penetrating the agar phantom. The electrode was lowered until the targeted depth (i.e., approximately 5, 10, 15, and 20 mm) was reached. Images of the insertion trajectory were used to calculate the deflection or the radial length from the axis of insertion at the final depth of insertion. Deflection distance was computed as the distance of the electrode tip from axis of insertion in terms of radial lengths as determined from two imaging planes. We fitted a linear regression line to the measured deflection as a function of insertion depth (**Fig. 4a**). For blunt-cut CFETs, the estimated beta coefficient for insertion depth was 0.40 with an intercept term of –0.79. The model fit r-squared was 0.327, with an f-statistic of 19.9 versus a degenerate constant model (p = 6.2e-05); there were 2 degrees of freedom for this linear model with 41 error degrees of freedom. For flame-etched CFETs, the estimated beta coefficient for insertion depth was 0.11 with an intercept term of 0.187. The model fit r-squared was 0.385, with an f-statistic of 11.9 versus a degenerate constant model (p = 0.002); there were 2 degrees of freedom for this linear model with 19 error degrees of freedom. These fits were then used to predict the deflection of the CFETs at a fixed insertion depth (e.g., 5 mm and 18.5 mm) for both cut-types of CFETs, as the insertion-depth was variable across tests. Areal coverage was also estimated from these fits. These steps are also described at protocols.io (dx.doi.org/10.17504/protocols.io.kxygx9b5zg8j/v1).

### Histology and Retention of Implanted Electrodes in Fixed Tissue

Rats were deeply anesthetized with Euthasol (pentobarbital sodium and phenytoin sodium from Virbac AH Inc.), then transcardially perfused with 0.9% sodium chloride (saline) solution followed by 4% paraformaldehyde in 0.1 M phosphate buffer (PB). The implanted electrodes were cut just above the skull and below the copper wire bundle component of the device with a rotary saw (Dremel) to retain the flexible Py-CF portion of the device in the fixed brain tissue. Brains were removed from the skull, post-fixed for 24 hours in the same fixative solution and stored in 25% glycerol with 0.075% sodium azide in 0.1 M PB overnight or until cutting. To prepare sections, brains were frozen on dry ice then cut in the transverse plane on a freezing microtome with a microtome knife (C. L. Sturkey, Inc. #K185A). Sections were cut sequentially, with four sections at 25-μm then one section at 100-μm, repeated until the end of the frozen block of brain. The 100-μm sections were immediately mounted onto glass slides and air dried, whereas the 25-μm sections were stored in 0.1% sodium azide in 0.1 M PB at 4°C until use. This protocol is also described at protocols.io (dx.doi.org/10.17504/protocols.io.6qpvr4nm3gmk/v1).

### Immunohistochemistry

For the immunofluorescent staining, free floating 25-μm sections from rat A were rinsed three times for 5 min each with 0.01 M phosphate buffered saline with 0.2% Triton X-100 (PBS-Tx), and then blocked for 60 min in TSA Blocking reagent (Akoya science FP1012) diluted in PBS-Tx (TSA block) on a shaker. Sections were left shaking for one night at 4°C in primary antibodies diluted in TSA block. Primary antibody and concentration used was: rabbit anti-MOR1(Abcam Ab134054, 1:500). Sections were rinsed three times for 5 min each with PBS-Tx and then were incubated in secondary antibodies diluted in TSA block for 2 hours on the shaker at room temperature. Secondary antibodies and concentrations used were: goat anti-rat conjugated with AlexaFluor 546 (Invitrogen A-11081, 1:300), and goat anti-rabbit conjugated with AlexaFluor 647 (Invitrogen A-21245, 1:300). Sections were rinsed three times for 5 min in 0.1 M PB, then incubated for 2 min in DAPI (Invitrogen 62248, 1:1000) diluted in PBS. After rinsing three times for 5 min in 0.1 M PB, all sections were mounted onto glass slides and cover slipped with ProLong Gold antifade reagent (Invitrogen, P36930). Slides were covered with aluminum foil and stored at 4°C until imaging. 100-μm sections on the glass slides from rat A were cover slipped without any staining with ProLong Gold antifade reagent.

The 100-μm sections on the glass slides from rat B were rinsed three times for 5 min each with 0.01 M PBS-Tx and were blocked endogenous peroxidase activity with 3% H_2_O_2_ in PBS-Tx. After three times washing with PBS-Tx, slides were blocked in TSA block in the humidity-controlled chamber. After removing the excess liquid from slides, rabbit anti-MOR1 (Abcam Ab134054, 1:500) antibody diluted in TSA block was applied on the slides and slides were left for one night at 4°C in a humidity-controlled chamber. Slides were then rinsed three times for 5 min each with PBS-Tx and were incubated with goat anti-rabbit antibody conjugated with polymer HRP (Thermofisher B40962) for 45 minutes at room temperature. After rinsing three times for 5 min each with PBS-Tx, slides were incubated with TSA-Plus Fluorescein (PerkinElemer NEL745001KT) diluted at 1:100 in 1X Plus Amplification Diluent for 15 minutes at room temperature. Slides were rinsed three times for 5 min in 0.1 M PB, then incubated for 2 min in DAPI (1:1000, Invitrogen 62248) diluted in PBS. After rinsing three times for 5 min in 0.1 M PB, slides were air dried, and then were cover slipped with ProLong Gold antifade reagent. Slides were covered with aluminum foil and stored at 4°C until imaging was done.

The TissueFAXS Whole Slide Scanning System (TissueGnostics) with x20 objective lens was used to obtain tiled-images from rat A. Cameras equipped with this system are Baumer HXG40c (HX series) CMOS camera 16 bit (2048 × 2048) for brightfield imaging and Hamamatsu Orca Flash 4.0 V2 cooled digital CMOS camera C11440-22CU for fluorescence imaging. Brightfield images were obtained from 100-μm sections to see carbon fibers left in the brains and fluorescence images were obtained from 25-μm sections to visualize MOR1 enriched striosome and endogenous rat IgG. These steps are also described at protocols.io (dx.doi.org/10.17504/protocols.io.n92ldpx78l5b/v1).

## Data Availability

All source data are available at zenodo.org (10.5281/zenodo.7818783). All protocols are available at protocols.io.

## Code Availability

MATLAB code used to analyze data are available at the GitHub repository (https://github.com/hschwerdt/multifscv).

## Acknowledgment

The authors thank Mr. H.F. Hall and Drs. D. Hu and Y. Kubota (Graybiel Laboratory, Massachusetts Institute of Technology), and Dr. Esta Abelev (Nanoscale Fabrication and Characterization Facility, University of Pittsburgh) for help with fabrication and surgical procedures, research insight, and manuscript preparation. This work was performed, in part, at the Nanoscale Fabrication and Characterization Facility, a laboratory of the Gertrude E. and John M. Petersen Institute of NanoScience and Engineering, housed at the University of Pittsburgh. This work was supported by NIH NINDS (R00 NS107639 to H.N.S.), BBRF NARSAD Young Investigator Grant (28690 to H.N.S.), the Michael J. Fox Foundation for Parkinson’s Research (MJFF) and the Aligning Science Across Parkinson’s (ASAP) initiative (ASAP-020-519 to H.N.S.), NIH NIMH (R01 MH060379 to A.M.G.), the Saks Kavanaugh Foundation (to A.M.G.), and the William N. and Bernice E. Bumpus Foundation (to A.M.G.). MJFF administers the ASAP-020-519 on behalf of ASAP and itself.

## Author Contributions

H.N.S., M.X., and B.N.A. designed experiments. H.N.S., A.M.G., and M.J.C. guided experimental design. M.X., B.N.A., K.B., and H.N.S. fabricated devices and performed *in vivo* and *in vitro* procedures. J.C., U.A., N.K., and S.Z., performed *in vitro* experiments. T.Y. performed histology and tissue-retainment experiments. H.N.S., M.X., and A.M.G. wrote manuscript with comments from all other authors.

